# Plasma Membrane Calcium ATPase (PMCA) Regulates Stoichiometry of CD4^+^ T-cell Compartments

**DOI:** 10.1101/2021.03.18.435950

**Authors:** Maylin Merino-Wong, Barbara A. Niemeyer, Dalia Alansary

**Author notes:** Corresponding author: Molecular Biophysics, Center for Integrative Physiology and Molecular Medicine (CIPMM), Building 48, School of Medicine, Saarland University, 66421 Homburg, Germany.

## Abstract

Immune responses involve mobilization of T cells within naïve and memory compartments. Tightly regulated Ca^2+^ levels are essential for balanced immune outcomes. How Ca^2+^ contributes to regulating compartment stoichiometry is unknown. Here, we show that plasma membrane Ca^2+^ ATPase 4 (PMCA4) is differentially expressed in human CD4^+^ T compartments yielding distinct store operated Ca^2+^ entry (SOCE) profiles. Modulation of PMCA4 yielded a more prominent increase of SOCE in memory than in naïve CD4^+^ T cell. Interestingly, downregulation of PMCA4 reduced the effector compartment fraction and led to accumulation of cells in the naïve compartment. *In silico* analysis and chromatin immunoprecipitation unraveled Ying Yang 1 (YY1) as a transcription factor regulating PMCA4 expression. Analyses of PMCA and YY1 expression patterns following activation and of PMCA promoter activity following downregulation of YY1 highlight repressive role of YY1 on PMCA expression. Our findings show that under the transcriptional control by YY1, PMCA4 adapts Ca^2+^ levels to cellular requirements during effector and quiescent phases and thereby represent a potential target to intervene with the outcome of the immune response.

## Introduction

The spatiotemporal characteristics of Ca^2+^ signals in T cells tightly control the outcome of an immune response as well as the fate of the immune cells involved. A local or global rise of intracellular [Ca^2+^]_i_ is achieved by activation of Ca^2+^ channels or by mobilization from intracellular stores. Once the Ca^2+^ signal is conducted, extrusion pumps restore the resting concentration. Alternatively, resting concentration is restored by Ca^2+^ uptake into intracellular stores: mitochondria or endoplasmic reticulum (ER).

Plasma membrane Ca^2+^ ATPase (PMCA) is the major extrusion pump playing a pivotal role in regulation of Ca^2+^ dynamics during activation of T cells (Bautista, Hoth et al., 2002, Bautista & Lewis, 2004, Lewis, 2001, Samakai, Hooper et al., 2016) in co-operation with sodium calcium exchanger (NCX) (Kiang et al., 2003) and the sarco-endoplasmic reticulum Ca^2+^ ATPase (SERCA) (Launay, Bobe et al., 1997). In addition to its global role, PMCA is able to generate local gradient in its vicinity thereby influencing Ca^2+^ dependent interaction partners (Stafford, Wilson et al., 2017). Furthermore, PMCA interacts with the inhibitory domain of CD147 in a Ca^2+^ independent manner which is necessary for CD147-mediated inhibition of IL-2 production (Supper et al., 2016). PMCA is a type-P ATPase of the class PIIB (Brini, Cali et al., 2013). In mammals PMCA contains a C-terminal auto-inhibitory domain, the inhibition by which is alleviated by binding to calmodulin or acidic phospholipids which can also bind to a basic domain in the first cytosolic loop (reviewed in (Gong, Chi et al., 2018)). There are four genes expressing PMCA (*ATP2B1-4*) with two possible splice sites resulting in a sum of about 30 variants belonging to 4 isoforms (Strehler & Zacharias, 2001). All isoforms of PMCA have Ca^2+^ dependent activity with PMCA4b being the slowest isoform despite a higher affinity to Ca^2+^/calmodulin than PMCA4a (Caride, Filoteo et al., 2001). The immunoglobulin proteins basigin and neuroplastin were identified as obligatory subunits of PMCA isoforms (Korthals, Langnaese et al., 2017, Schmidt, Kollewe et al., 2017). Co-assembly into a complex with either protein is necessary for stable PMCA surface localization in rat brain derived neurons (Schmidt et al., 2017) but also murine cells lacking neuroplastin showed reduced expression of PMCA (Korthals et al., 2017).

The past four decades witnessed development of models describing the differentiation of naïve T cells following antigen encounter into different compartments of effector and memory cells. These models are based on flow cytometric phenotypic analysis and their outcome provides the basis for a nomenclature of subpopulations and describes function and persistence of populations arising during antigen encounter and clearance (for comprehensive reviews see (Durek, Nordstrom et al., 2016, Farber, Yudanin et al., 2014, Sallusto & Lanzavecchia, 2009)). There are three main models describing memory cell generation. The first is based on epigenomic profiling and proposes linear development of naïve into stem central memory (TSCM) then CM, followed by effector memory (EM) and eventually effector (E) cells (Durek et al., 2016). The bifurcative model suggests that naïve cells asymmetrically divide to directly develop into E cells and in parallel an EM that gives rise to a CM lineage. The third model overlaps with the linear model with the effector cells retaining ability for self-renewal and hence the name of the model (Ahmed, Bevan et al., 2009). Despite the intensive efforts, the mechanisms regulating naïve-to-memory cell transition are not entirely understood. One important aspect of this transition is the transcriptional reprogramming of the cells (Christie & Zhu, 2014, Schoettler, Hrusch et al., 2019). Through iterative interactions, transcription factors selectively combine genes to be expressed to define cytokine production and survival potential profiles characteristic of cell populations. Ca^2+^ dependence is a common feature of many of the transcription factors involved in T cell differentiation such as NFAT (Jain, McCaffrey et al., 1993, Rao, Luo et al., 1997), CREB (Mayer & Thiel, 2009) and the transcription repressor DREAM (Savignac, Mellstrom et al., 2007). The main pathway for Ca^2+^ influx in T cells is the store operated Ca^2+^ entry (SOCE) constituted by the channel forming ORAI proteins (Vig, Peinelt et al., 2006, Zhang, Yeromin et al., 2006) and the activator ER-resident Ca^2+^ sensor STIM proteins (Liou, Kim et al., 2005, Roos, DiGregorio et al., 2005, Zhang, Yu et al., 2005). While Ca^2+^ levels influence the activity of transcription factors, conversely, the expression of SOCE components (Ritchie, Yue et al., 2010) and also expression of proteins involved in Ca^2+^ homeostasis is subjected to transcriptional control (Ritchie, Zhou et al., 2011). Transcriptional control of PMCA is, however, poorly understood and whether this control is altered during naïve-to-memory cell transition is yet to be explored. Therefore, we set out in the current study to investigate how PMCA alters Ca^2+^ signals in T cells to promote activation and differentiation; and to gain insight into mechanisms regulating PMCA expression during T cell activation.

## Results

### Differential expression of PMCA4b results in distinct SOCE profiles in CD4^+^ naïve and memory cells

In our previous work we characterized the SOCE profiles of antiCD3/CD28-activated cells, *in vitro* polarized into CD4^+^ T cell subtypes (Th1, Th2, Th17 and Treg) and observed that the control naïve and memory cells, that were activated but cultured in non-polarizing conditions, showed differential SOCE profiles with concomitant upregulation of PMCA in the activated memory cells (Kircher, Merino-Wong et al., 2018). Therefore, we set out in the current work to investigate whether in resting cells PMCA plays a differential role in Ca^2+^ homeostasis and thus regulates the differentiation process of human CD4^+^ T cells. First, we measured SOCE profiles in resting naïve (CD4^+^CD127^high^CD25^-^CD45RO^-^) and memory (CD4^+^CD127^high^CD25^-^CD45RO^+^) CD4^+^ T cells. We isolated the desired populations from human peripheral blood mononuclear cells (PBMCs) using the fluorescence activated cell sorting (FACSorting) strategy shown (Fig. S1) where we excluded regulatory cells (Treg, CD4^+^CD127^low^CD25^+^) and sorted the desired populations from conventional CD4^+^ T cells (CD4^+^CD127^high^CD25^-^). One day later, we measured changes of intracellular Ca^2+^ concentration following thapsigargine (Tg)-induced store depletion. The resulting SOCE profiles of naïve and memory human CD4^+^ cells were significantly distinct. Naïve cells showed a higher peak and plateau of [Ca^2+^]_i_ than memory cells (Fig. 1A, B). The observation that the peak [Ca^2+^]_i_ of memory cells was on reduced by 13.5 % while the steady state plateau [Ca^2+^]_i_ and the fraction of retained Ca^2+^ (Plateau/Peak) were reduced by more than 40% each, compared to naïve cells indicates that Ca^2+^ signals in memory cells exhibit faster decay kinetics. To determine whether the difference in naïve and memory profiles is due to differential expression of Ca^2+^ influx and/or extrusion mechanisms we analyzed the expression of the relevant genes. First, we analyzed the expression levels of *ORAI1-3* and *STIM1-2*, the main constituents of Ca^2+^ influx pathways in T cells. Analysis of mRNA (Fig. 1C) and protein (Fig. 1D, E) levels showed that both naïve and memory CD4^+^ cells express comparable levels of SOCE components. The specificity of the antibodies was validated by cells where *ORAI1 (Alansary, Peckys et al., 2020), STIM1* or *STIM2* (Ramesh, Jarzembowski et al., 2021) were deleted by CRISPR-Cas9 mediated gene targeting. Using commercially available antibodies for ORAI2 and ORAI3 proteins we were unable to reliably detect endogenous proteins. These results indicate that reduced [Ca^2+^]_i_ is likely not explained by reduced Ca^2+^ influx through ORAI channels and imply involvement of Ca^2+^ efflux mechanisms. Analysis of expressed isoform of plasma membrane Ca^2+^ dependent ATPase (PMCA, *ATP2B1-4*) revealed that the main isoforms expressed in CD4^+^ cells are PMCA1 (*ATP2B1*) and the more predominant PMCA4 (*ATP2B4*) in agreement with (Caride et al., 2001) while isoforms PMCA2 (*ATP2B2*) and PMCA3 (*ATP2B3*) were not detected (Fig. 1F). Furthermore, using splice-specific primers, mRNA analysis showed that PMCA4b (*ATP2B4b*) is the main variant in CD4^+^ T cells (Fig. 1G). Interestingly, memory cells showed a significantly higher expression level of PMCA4b (*ATP2B4b*) on both mRNA level (Fig. 1H) and protein (Fig. 1I, J) level, using an antibody with tested specificity ((Ho, Pang et al., 2015) and Fig. S2B). Moreover, we analyzed Ca^2+^ efflux rates in presence (Fig. 1K) and after removal (Fig. 1L) of extracellular Ca^2+^, taking into consideration the Ca^2+^ dependence of PMCA activity (Bautista et al., 2002, Bautista & Lewis, 2004). By binning the cells according to [Ca^2+^]_i_ allowed comparison of cells with similar [Ca^2+^]_i_ (iso-cells) and revealed that memory cells extrude Ca^2+^ with higher rates compared to naïve cells (Fig. 1K, L) independent of the cytosolic [Ca^2+^]. Significant differences are observed at physiologically relevant concentrations but not at higher concentrations probably due to exceeding the pumping capacity of PMCA but also due to the small fraction of cells exhibiting this concentration which hampers statistical significance.

**Fig. 1.**
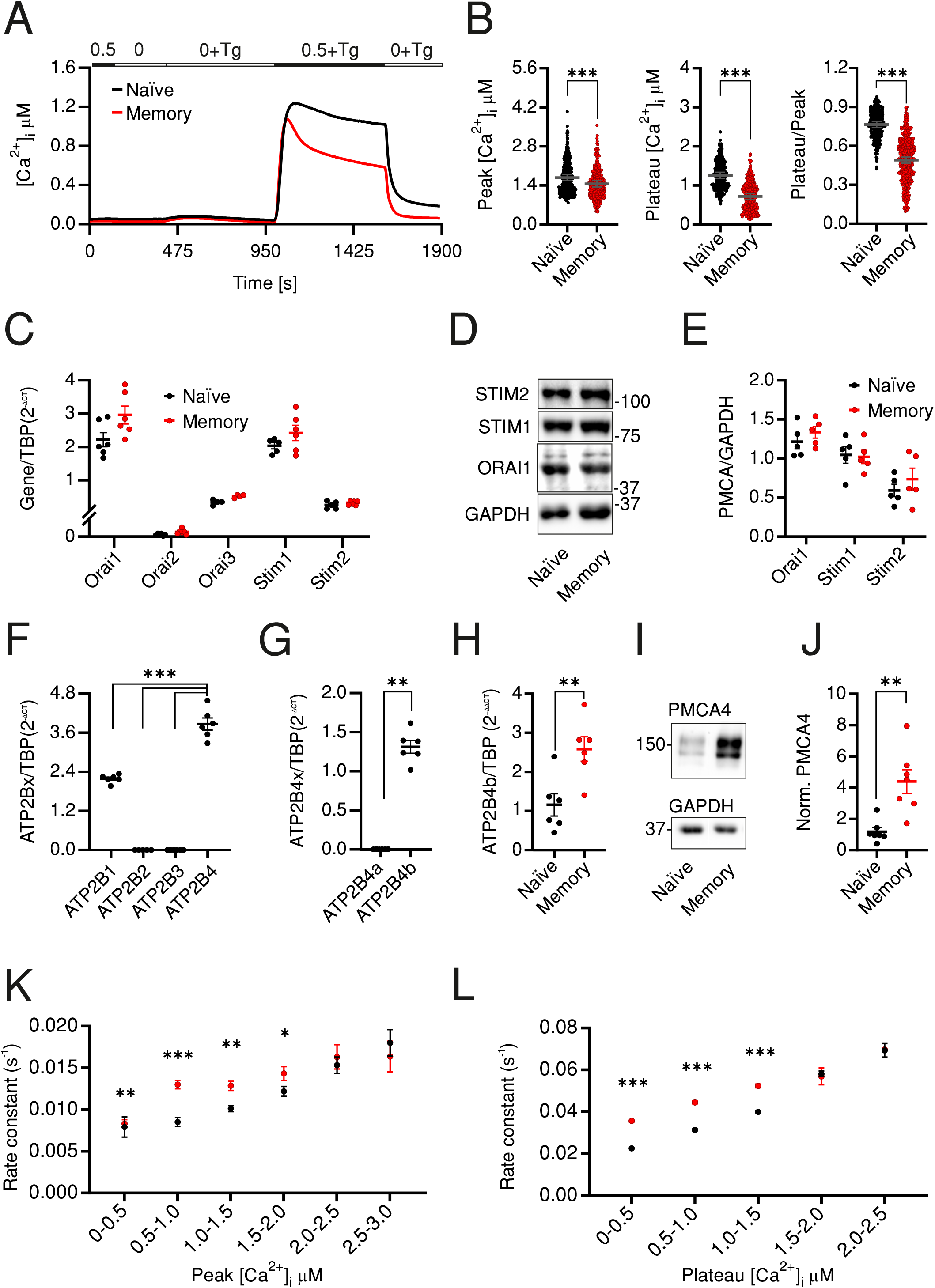
CD4 naïve and memory cells have distinct SOCE profiles due to differential expression of PMCA4b. (A) Average traces showing changes of [Ca^2+^]_i_ over time in response to changes of extracellular solutions containing the shown [Ca^2+^]_o_ indicated in the bar above the traces. SOCE was measured in human naïve (CD4^+^CD127^high^CD25^-^CD45RO^-^, black) or memory (CD4^+^CD127^high^CD25^-^CD45RO^+^, red) cells isolated from PBMCs. (B) Bar graphs showing analyzed parameters of SOCE measured in (A). (C) The relative expression of *ORAI1-3* and *STIM1-2* mRNA in cells measured (A). (D) Representative of 5 independent Western blots (WB) and corresponding quantification (E) analyzing expression of ORAI1, STIM1 and STIM2 in cells measured in (A). (F-H) Quantitative RT-PCR analysis showing (F) the relative expression of the four isoforms of PMCA (G) relative expression of the splice variants 4a and 4b in total CD4^+^ cells, and (H) relative expression of PMCA4b in naïve (black) and memory (red) cells isolated as in (A) from human PBMCs. (I) Representative of 5 independent WB and corresponding quantification (J) showing expression of PMCA4 in cells measured in (A, H). (K) Analysis of Ca^2+^ efflux rates in presence of [Ca^2+^]_o_ in cells measured in (A) as function of peak [Ca^2+^]_i_ (L) Analysis of Ca^2+^ efflux rates after removal of [Ca^2+^]_o_ in cells measured in (A) as function of plateau [Ca^2+^]_i_. Data represent average ± s.e.m obtained from 3257 and 2644 cells measured in 22 and 23 independent experiments (shown dots in A, B), 5-6 independent experiments (C-J). Asterisks indicate significance at ** p<0.01, *** p<0.001 using unpaired two-tailed Student t-test (B), Mann-Whitney test (G, J) or Kruskall-Wallis one-way analysis of variance (ANOVA) (C-F)

### Pharmacological inhibition or downregulation of PMCA reverses SOCE phenotypes of memory cells

To directly test whether PMCA has a differential role in Ca^2+^ homeostasis in resting CD4^+^ T cell subsets, we tested the effect of pharmacological inhibition of PMCA by caloxin1C2 (C1C2) (Pande, Szewczyk et al., 2011) on SOCE profiles of naïve and memory cells. Treatment with 5 μM or 20 μM caloxin1C2 only mildly altered SOCE profile of the naïve cells resulting in no or a slight increase that did not exceed 14% of the tested parameters (Fig. 2A, B). On the other hand, memory cells treated acutely with the PMCA blocker showed a significantly increased SOCE resulting in a 58% increase of the plateau and 47% increase of the fraction of retained Ca^2+^ compared to untreated cells (Fig. 2C, D). Noteworthy is that the application of 20 μM, a concentration shown to block all PMCA isoforms (Pande et al., 2011), did not result in additional effects on the plateau levels compared to application of the PMCA4-specific inhibitory concentration 5 μM (Fig. 2C, D compare blue to grey traces and dots) indicating that PMCA4 is indeed the main functional isoform in T cells and in agreement with the expression analyses (Fig. 1F-J).

**Fig. 2.**
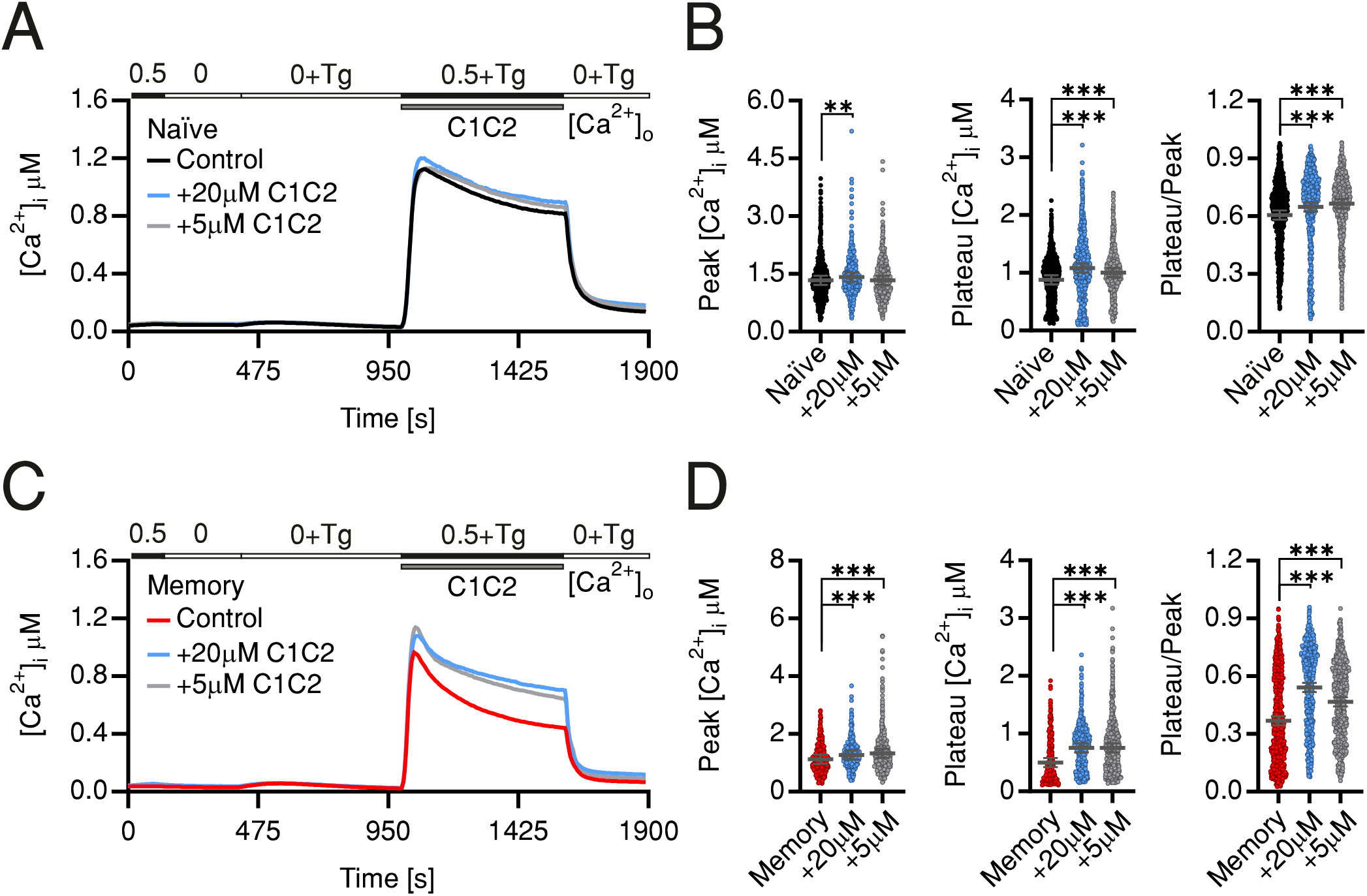
Pharmacological inhibition of PMCA reverts SOCE phenotypes of memory cells. Average traces showing changes of [Ca^2+^]_i_ over time in response to changes of extracellular solutions containing the shown [Ca^2+^]_o_ indicated in the bar above the traces. SOCE was measured in human naïve (A, black) or memory (C, red) control cells or corresponding cells treated with 20 μM Caloxin1C2 (blue), or with 5 μM Caloxin1C2 (grey). (B, D) Bar graphs showing analyzed parameters of SOCE measured in (A, C), respectively. Data represent average ± s.e.m obtained from 976 to 2659 cells, measured in 8-10 independent experiments. Asterisks indicate significance at ** p<0.01, *** p<0.001 using Kruskall-Wallis one-way analysis of variance (ANOVA).

In a second approach we transfected naïve or memory CD4^+^ T cells using an *ATP2B4* specific siRNA (siPMCA) that resulted in significant downregulation on RNA (Fig. S2A) and protein levels (Fig. S2B). Similar to the pharmacological manipulation by C1C2, downregulation of PMCA4 led to no or subtle alteration of analyzed parameters of SOCE profile of the naïve cells (Fig. 3A, B). Memory cells treated with siPMCA showed a 15 % increased influx peak while the plateau was increased by 49 % compared to memory cells transfected with non-silencing RNA (ns) resulting in significant increase in fraction of Ca^2+^ retained in the cells (Plateau/Peak ratio) (Fig. 3C, D). These results indicate a differential role of PMCA in Ca^2+^ homeostasis of naïve and memory cells.

**Fig. 3.**
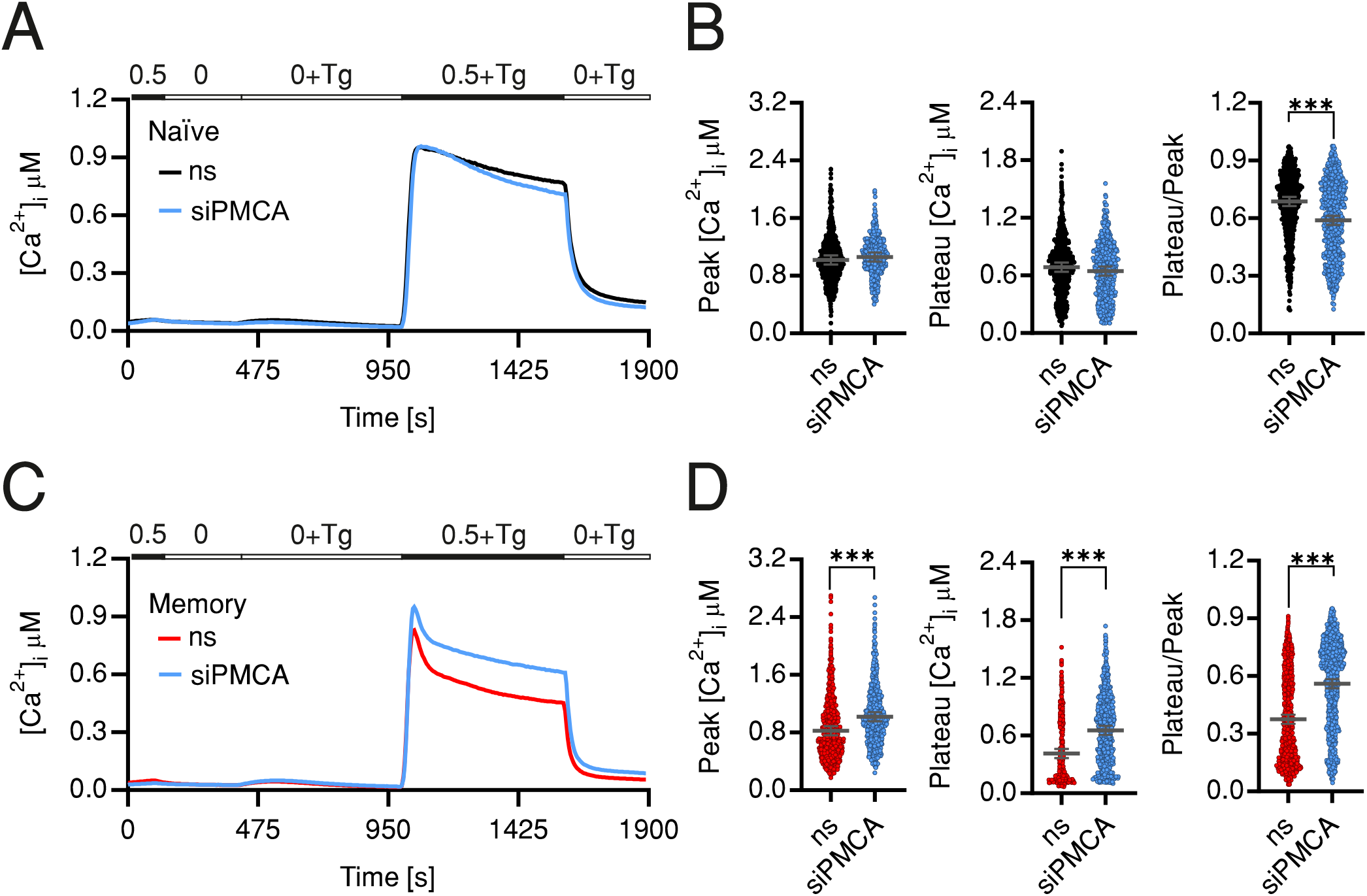
Downregulation of PMCA reverts SOCE phenotypes of memory cells. Average traces showing changes of [Ca^2+^]_i_ over time in response to changes of extracellular solutions as shown in the bar above the traces. SOCE was measured in human naïve (A, ns, black) or memory (C, ns, red) cells transfected with non-silencing RNA or transfected with PMCA4 targeting siRNA (siPMCA, blue). (B, D) Bar graphs showing analyzed parameters of SOCE measured in (A, C). Data represent average ± s.e.m obtained from 1163 to 1280 cells, measured in 7-10 independent experiments (shown dots). Asterisks indicate significance at ** p<0.01, *** p<0.001 using Kruskall-Wallis one-way analysis of variance (ANOVA).

### In vivo developed effector and memory CD4^+^ cellular compartments differentially express PMCA4 and exhibit corresponding SOCE profiles

The findings presented so far compared the Ca^2+^ signals and expression profiles of naïve and memory cells. Memory cells (CD4^+^CD45RO^+^), however, represent a heterologous population that is commonly divided into different compartments depending on effector and proliferative capacities of the cells. Therefore, our next aim was to answer the question whether *in vivo* developed memory compartments show distinct PMCA expression profiles or do they homogenously share higher expression levels of PMCA. To address this question, we FACSorted CD4^+^ T cell compartments from human PBMCs based on the surface markers (CD45RO/ CCR7) as illustrated in the in (Fig. 4A). Purity of the sorted populations was tested by post-sorting staining (Fig 4B). Cells that were CCR7^+^CD45RO^-^ CD45RA^+^ are defined as naïve cells (N), CCR7^+^CD45RO^+^ CD45RA^-^ as central memory cells (CM), CCR7^-^ CD45RO^+^ CD45RA^-^ effector memory (EM) and CCR7^-^ CD45RO^-^ CD45RA^+^ as terminally differentiated effector memory cells (EMRA) (Mahnke, Brodie et al., 2013, Sallusto, Lenig et al., 1999, Tian, Babor et al., 2017). T_EMRA_ cells is a poorly characterized population of T cells expressing CD45RA but commonly used due to sharing functional properties with the short lived effector cells (Durek et al., 2016, Henson, Riddell et al., 2012, Sallusto, Geginat et al., 2004). For further characterization, we measured the expression of IFNγ which showed the expected pattern where the EMRA and EM populations had higher expression of IFNγ than the CM while it was lacking in naïve cells (Fig. 4C).

**Fig. 4.**
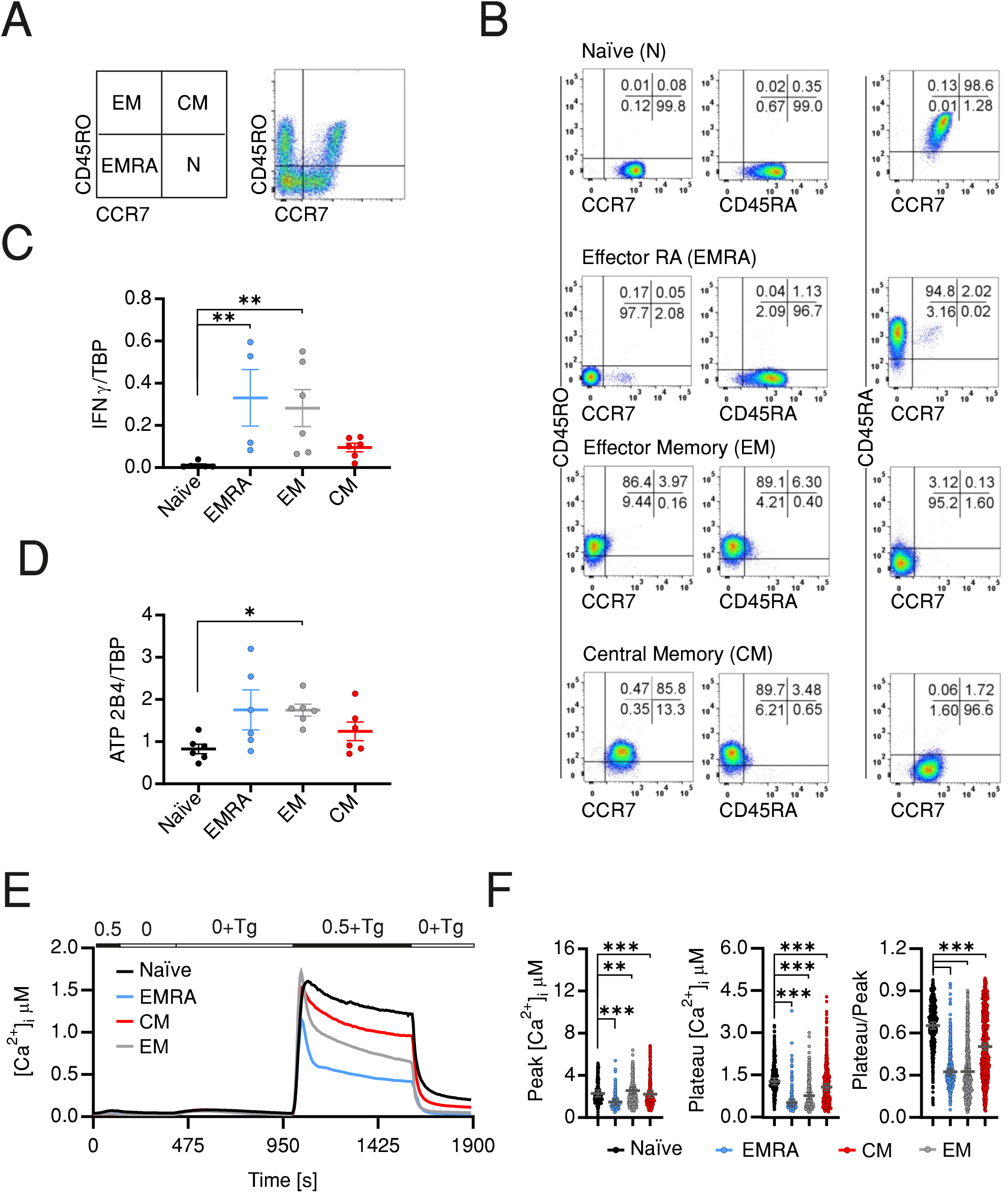
In vivo CD4^+^ effector and memory compartments differentially express PMCA and exhibit corresponding SOCE profiles. (A) Representative images showing sorting strategy of CD4^+^ cells into the different compartments using the shown surface markers and according to the shown scheme. (B) Representative images obtained by post-sorting flow cytometric analysis of the sorted cells. Quantitative RT-PCR analysis showing relative expression of *IFNG* (C) and PMCA4 (*ATP2B4*) (D) relative to TBP, in CD4^+^ compartments isolated as in (A). (E) Average traces showing changes of [Ca^2+^]_i_ over time in response to changes of extracellular solutions as shown in the bar above the traces, analyzed in CD4^+^ T cell compartments isolated as in (A). (F) Bar graphs showing analyzed parameters of SOCE measured in (E). Data represent average ± s.e.m obtained from 586-1084 cells measured in 6 independent experiments. Asterisks indicate significance at * p<0.05, **p<0.01 and *** p<0.001 using Kruskall-Wallis one-way analysis of variance (ANOVA)

Interestingly, analysis of PMCA expression in the sorted populations revealed differential expression profiles: with the terminally differentiated effector (EMRA) and EM cells showing the higher expression than CM and naïve cells (Fig. 4D). Finally, we tested whether SOCE profiles of the *in vivo* differentiated cellular compartments recapitulate the differentially expressed PMCA levels. Indeed, measurements of Tg-induced Ca^2+^ influx showed that CD4^+^ compartments have distinct SOCE profiles with comparable Ca^2+^ peaks (Fig. 4E, F) but distinct increase in decay rates resulting in lowest plateau [Ca^2+^]_i_ and retained Ca^2+^ fractions for the effector cells (EMRA and EM), intermediate levels for CM cells while the naïve cells have the highest plateau [Ca^2+^]_i_ and retained Ca^2+^ fractions (Fig. 4E, F) in line with the observed PMCA expression levels (Fig. 4D). These differences are underlined by parallel differences in the efflux rates in presence of (Fig. S2C) and after removal (Fig. S2D) of extracellular Ca^2+^ indicating that the differences in the plateau and retained Ca^2+^ fractions are indeed due to differential expression and function of PMCA.

### PMCA4 regulates compartment stoichiometry upon activation

The outcome of an immune response depends on the dynamics of mobilization of T cells within naïve and memory compartments. Furthermore, our findings show that CD4^+^ cellular compartments exhibit differential SOCE profiles and PMCA expression levels. Therefore, we sought to address if and how regulation of [Ca^2+^]_i_ by PMCA4 contributes to fate-decision making concerning compartment distribution of CD4^+^ T cells upon activation. To this end, we isolated naïve CD4^+^ T cells from PBMCs, to exclude effects of previous antigen exposure, and on the following day transfected naïve cells with non-silencing RNA (ns) or with siRNA targeting *ATP2B4* (siPMCA). Three days later, naïve cells were replenished with fresh siRNA, subjected to stimulation using antiCD3/CD28 coated beads and monitored for compartment distribution 24 h later. The experimental design and the surface markers used to assign activated cells into different compartments are shown in figure (5A). Down regulation of *ATP2B4* did not alter stoichiometry of cellular compartments as defined by CD45RO or CD45RA and CCR7 (Fig. S2E, F). Because, classically, T cell compartments are defined based on CD45RO, CD45RA, CCR7 and CD62L (Mahnke et al., 2013, Sallusto et al., 1999, Tian et al., 2017) and CD62L is commonly used an alternative marker for CCR7 for compartment definition (Eichmann, Baptista et al., 2020, Priesner, Aleksandrova et al., 2016), we analyzed compartment stoichiometry based on CD45RO and CD62L expression. Interestingly, this analysis showed that downregulation of *ATP2B4* resulted in significant decrease of the fraction of cells in terminally differentiated effector memory CD45RA^+^ compartment T_EMRA_ (CD4^+^CD62^-^LCD45RO^-^) while the fraction of cells remaining in the naïve compartment (CD4^+^CD62L^+^CD45RO^-^) was significantly increased compared to cells treated with nonsilencing RNA (Fig. 4B, C). The effects on EM and CM compartments showed similar tendencies as effects observed on EMRA and naïve compartments, respectively, albeit not significant (Fig. 5B, C).

**Fig. 5.**
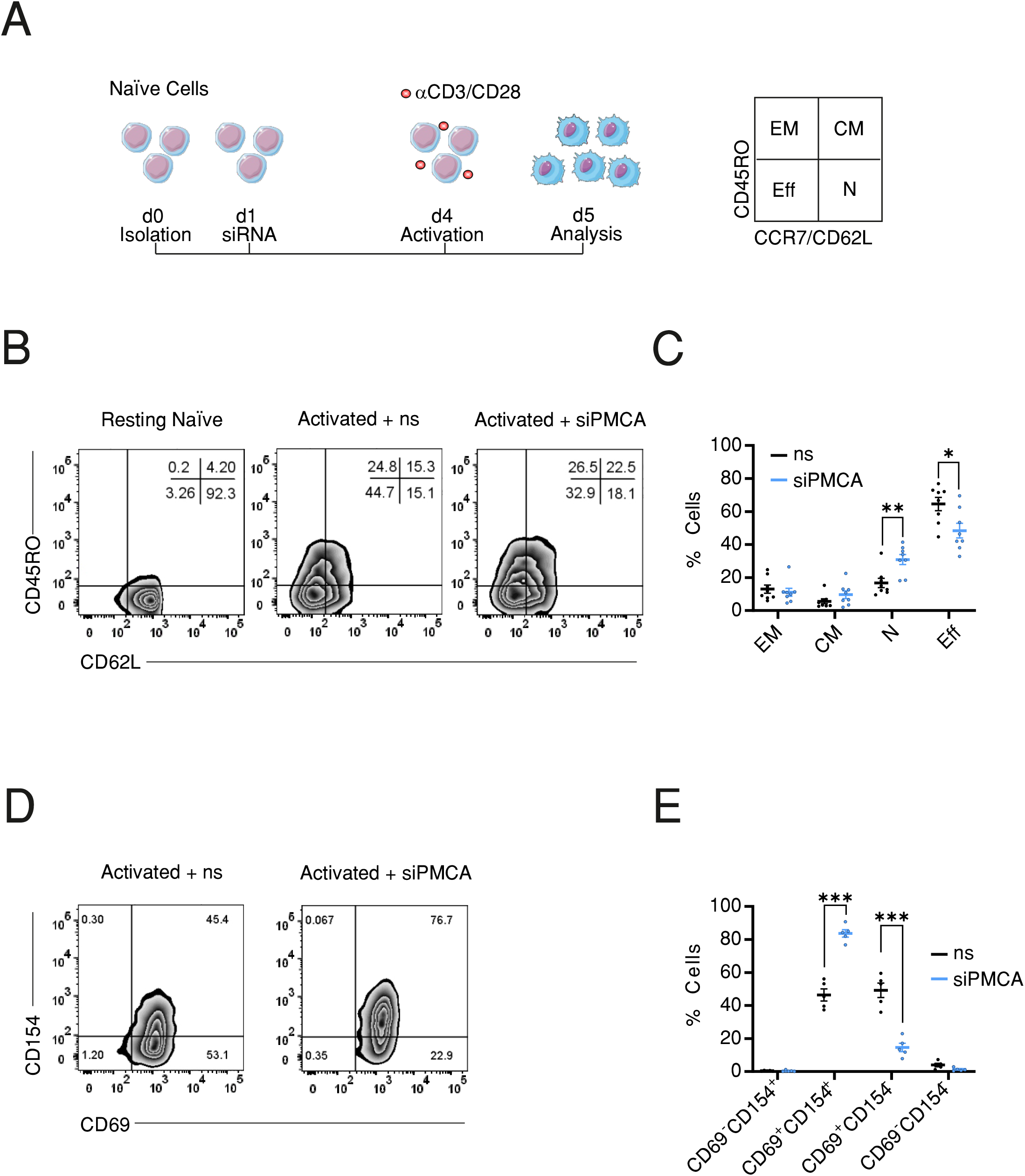
Downregulation of PMCA alters compartment stoichiometry upon activation of naïve CD4^+^ T cells. (A) Schematic representation of the experimental design: naïve cells were isolated from human PBMCs using FACSorting, transfected one day later with control non silencing RNA (ns) or with siRNA targeting PMCA4 (siPMCA), stimulated with antiCD3/CD28 coated beads 72 h following transfection and 24 h later analyzed for compartment distribution using flow cytometry and the shown surface markers. (B) Representative flow cytometric analysis images showing compartment distribution in resting naïve cells (left image) or naïve cells transfected with control non-silencing RNA (ns, middle image) or transfected with siRNA targeting PMCA4b (siPMCA, right image) then activated following time scheme shown in (A). (C) Frequency of cells (% Cells) measured in the EM (CD62L^-^CD45RO^+^), CM (CD62L^+^CD45RO^+^), N (CD62L^+^CD45RO^-^) and EMRA (CD62L^-^CD45RO^-^) compartments following treatment with non-silencing RNA (ns, black) or siRNA targeting PMCA4 (siPMCA, blue) according to time scheme shown in (A). (D) Representative flow cytometric analysis images showing expression of the activation markers CD154 and CD69 in cells treated as in (A, B). (D) Frequency of cells (% Cells) measured in (D) and expressing activation markers following treatment with non-silencing RNA (ns, black) or siRNA targeting PMCA4b (siPMCA, blue) according to time scheme shown in (A). Data represent average ± s.e.m obtained from 8 independent experiments. Asterisks indicate significance at * p<0.05, **p<0.01 using Kruskall-Wallis one-way analysis of variance (ANOVA).

To test whether downregulation of PMCA4 is accompanied by altered potential of naïve CD4^+^ cells to be activated, we measured the expression level of activation markers CD69 and CD154 under the same experimental conditions as above. Despite the reduced fraction of effector cells, treatment of naïve cells with siPMCA resulted in increased expression of activation marker CD154 (CD40L) (Fig. 5D, E) while expression of CD25 were not altered (Fig. S2G). These results imply that PMCA4 downregulation does not inhibit activation *per se* but rather regulates the compartment stoichiometry.

### Transcriptional control of PMCA results in biphasic expression following activation

The experiments described above provide evidence that PMCA4 expression level correlates with the cellular compartment of CD4^+^ T cells and contributes to regulation of compartment stoichiometry. To explore potential underlying mechanisms, we set out to investigate transcriptional regulation of PMCA expression. To this end, we explored potential transcription factor binding sites (TFBS) in the promoter region of PMCA (*ATP2B4*) using available ChIP seq information (Ambrosini, Dreos et al., 2016) combined with TFBS prediction data bases HOCOMOCO (Kulakovskiy, Vorontsov et al., 2018) and Promo 3 (Farre, Roset et al., 2003, Messeguer, Escudero et al., 2002). The resulting analysis (Fig. 6A) predicted 33 transcription factors (listed in supplementary Table. 1) that were detected in all three algorithms with the transcription factor YY1 having the highest predicted score. Consequently, we explored whether we can experimentally confirm binding of YY1 to the promoter region of PMCA. Because our preliminary experiments showed that a conventional protocol of chromatin immune precipitation (ChIP) experiment was prone to misleading positive results we applied a rigorous protocol that eliminates all unspecific binding using isotype control antibody bound beads in a “preclearing” step (Fig. 6B first lane). The efficiency of elimination of unspecific binding was demonstrated by applying the pre-cleared lysate to fresh isotype control antibody-bound beads (Fig. 6B second lane). Finally, the ability of YY1 to bind to the promoter region was tested by using a YY1-specific antibody to immunoprecipitate YY1 and the bound genomic DNA which was then amplified with specific primers, designed based on the predicted binding domains. In parallel we used an anti-NFAT antibody to serve as an additional control for the specificity of binding. Results of the ChIP experiments showed that YY1, but not NFAT, was able to bind the promoter region of PMCA4 as visualized on agarose gels (Fig. 6B). The identity of the amplicon was further confirmed by sequencing.

**Fig. 6.**
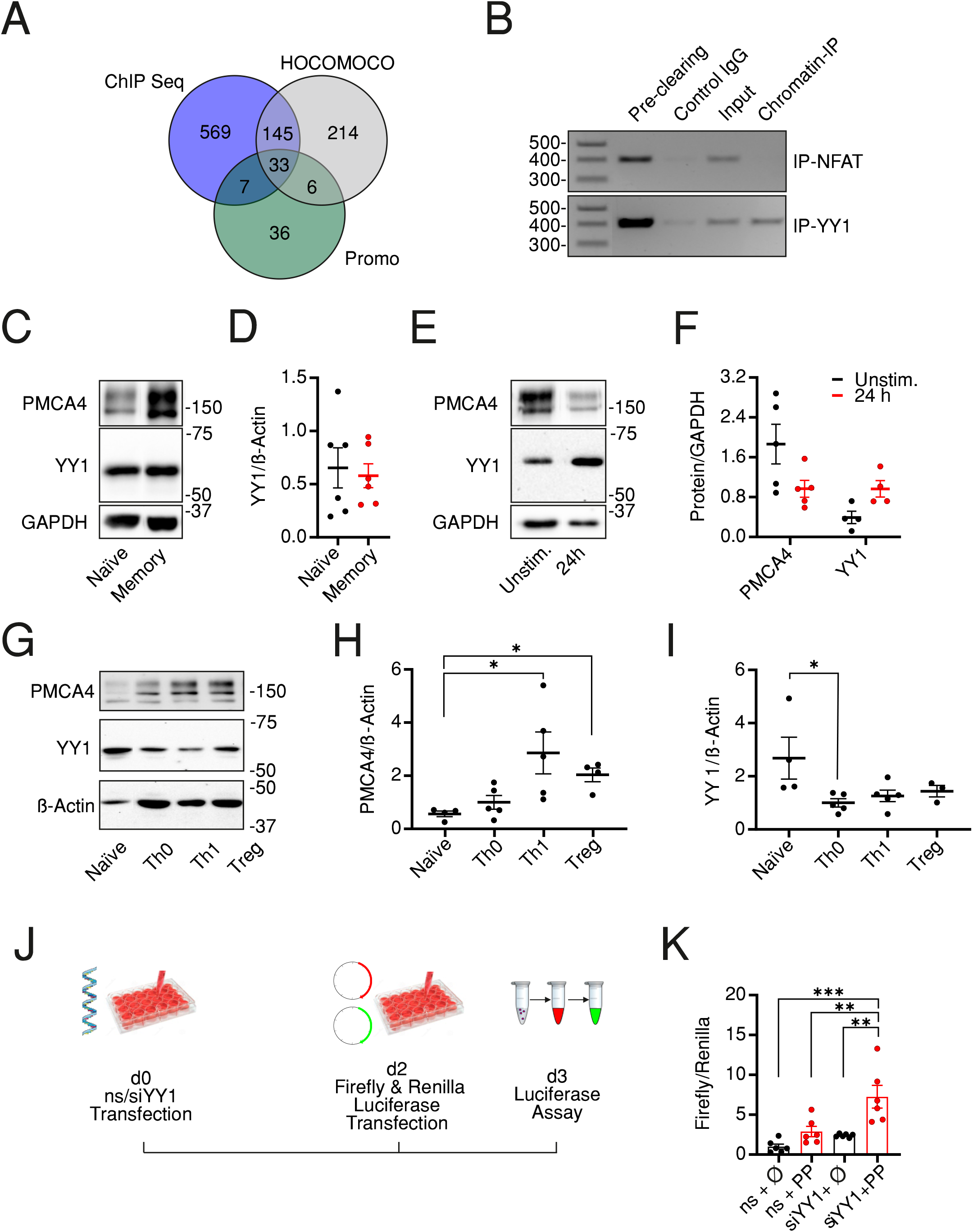
Transcriptional control of PMCA results in biphasic expression following activation. (A) Venn diagram showing the overlapping results obtained from three data bases (HOCOMOCCO, Promo V3 and Chip-seq) used to predict transcriptional factors binding to PMCA4 promoter region (see also materials and methods and supplementary info). (B) Agarose gel showing PCR amplicons obtained by using primers designed to amplify a 410 base pair-long fragment of the PMCA4b (*ATP2B4b*) promoter region. Poly chain reactions (PCRs) were performed using chromatin DNA purified from total lysate (Input) or after immune precipitation by pre-clearing beads, beads bound to IgG control antibodies, beads bound to anti-NFAT (upper row, Chromatin-IP) or to anti-YY1 (lower row, Chromatin-IP). (C) Representative of 5 WB and corresponding quantification (D) showing analysis of PMCA4 and YY1 in resting naïve or memory CD4^+^ T cells isolated as in Fig. 1. (E) Representative of 5 WB and corresponding quantification (F) showing analysis of PMCA4 and YY1 in total CD4^+^ T cells in resting conditions (Unstim.) or after 24 h stimulation with antiCD3/CD28 coated beads (24h). (G) Representative of 5 WB and corresponding quantification (H,I) showing analysis of PMCA4 (H) and YY1 (I) in naïve CD4^+^ T cells before (Naïve) and after 7d culture in control (Th0) or polarizing conditions into Th1 or regulatory (Treg) cells. (J) Schematic representation of experimental design of promoter assay: HEK293 cells were transfected with non-silencing RNA (ns) or with siRNA targeting YY1 (siYY1). On day 2, cells were transfected with plasmids expressing red firefly or green renilla luciferases and the dual luciferase assay was conducted on day 3. (K) Firefly luminescence normalized to Renilla luminescence in cells transfected as in (J) with firefly luciferase expressed in plasmid without promoter (ϕ) or with PMCA promoter (PP)

To investigate whether the levels of YY1 are altered in CD4^+^ cell compartments similar to PMCA, we analyzed expression levels in naïve and memory CD4^+^ cells. Surprisingly, expression level of YY1 were comparable in resting naïve and memory CD4^+^ cells (Fig. 7C, D). We hypothesized that this finding might be due to performing the analysis in resting cells where YY1 expression is adapted to steady state levels. To test this hypothesis, we stimulated total CD4^+^ cells for 24 h using antiCD3/CD28 coated beads and indeed we observed an increased level of YY1 concomitant to decreased expression of PMCA (Fig. 6E, F), indicating that YY1 represses PMCA levels upon stimulation, resulting in reduced Ca^2+^ clearance compared to unstimulated cells (Fig. S3A, B). Because PMCA levels were decreased after acute stimulation (Fig. 6E, F) contrary to the increased levels we observed in resting memory compared to naïve cells (Fig. 1H-J), we extended the expression analysis to examine whether YY1 alters PMCA4 expression in association with long term stimulation and full differentiation of T cells. To this end, we subjected the naïve cells to *in vitro* polarization protocol providing stimulatory, co-stimulatory and cytokine mediated signals necessary for differentiation of naïve cells into Th1 or Treg cells (Kircher et al., 2018). This experiment not only aims to monitor changes of PMCA and YY1 expression levels upon longer stimulation but also monitors correlation of PMCA expression to developing effector functions. The polarized cells showed the expected differential expression of *IFNG* (Fig. S3C) and *FOXP3* (Fig. S3D), the hallmarks of Th1 and Treg populations, respectively. Importantly, terminally differentiated effector cells (Th1, Treg) showed a significant upregulation of PMCA4 while the control cells which were stimulated in non-polarizing conditions (Th0) underwent milder upregulation of PMCA (Fig. 6G, H and Fig. S3E). Strikingly, levels of YY1 followed a reverse pattern: while YY1 underwent transient upregulation within 24 h of activation (Fig. 6E, F) YY1 expression decreased upon longer activation (Fig. 6G, I and Fig. S3F), in line with the equal levels analyzed in resting cells (Fig. 6C, D).

Finally, we aimed to confirm that altered expression of PMCA is due to direct regulation by YY1. To achieve this correlation, we constructed a plasmid expressing red firefly luciferase under transcriptional control of the identified PMCA promoter region where YY1 is predicted to bind (Fig 6A, B). To facilitate analysis of promoter activity independent of transfection efficiency, the red firefly luciferase activity was normalized to the co-transfected green renilla luciferase expressed under the transcriptional control of TK promoter. To address whether the PMCA promoter activity is directly regulated by YY1 we downregulated YY1 using siRNA (Fig. S3G, H) then conducted the dual luciferase assay (Fig. 6J). Measurements in (Fig. 6K) show that in cells transfected with non-silencing RNA (ns),, the PMCA promoter (ns + PP) was able to induced expression of the firefly luciferase compared to control cells (ns + ϕ). More importantly, the PMCA promoter activity is significantly increased following downregulation of YY1 (siYY1 + PP). Collectively, these results indicate that T cells adapt expression of PMCA4 during activation states under the transcriptional repressive control by YY1.

## Discussion

In the current work we investigated how Ca^2+^ signals are fine-tuned during activation of CD4^+^ T cells to regulate the stoichiometry of CD4^+^ T cell compartments and thus the outcome of an immune response. Using fura-2 Ca^2+^ imaging we show that quiescent memory cells accommodate a lower [Ca^2+^]_i_ and are able to clear the Ca^2+^ signals faster than their naïve counter parts (Fig. 1A, B). Comparable levels of expressed SOCE genes (Fig. 1C-E) but significantly more expressed PMCA4b (Fig. 1F-J) and higher efflux rates independent of [Ca^2+^]_i_ (Fig. 1K, L) present strong evidence that SOCE phenotype of memory cells is mostly a result of more efficient clearance machinery. Furthermore, analysis of the contribution of PMCA to Ca^2+^ homoeostasis in naïve and memory cells by applying an acute pharmacological inhibition approach (Fig. 2) as well as by directly downregulating *ATP2B4* (PMCA4) (Fig. 3) present evidence that clearly supports the hypothesis that PMCA4 play a functionally more significant role in Ca^2+^ clearance in memory compared to naïve CD4^+^ T cells. Further analysis of the CD4^+^ T cells subpopulations revealed that the quiescent cells, in the naïve and central memory compartments, express less PMCA and exhibit slower Ca^2+^ signal decay kinetics than compartments associated with more effector functions (EMRA and EM) (Fig. 4). *In vitro* generation of effector populations (Th1, Treg) able to express effector hallmarks (Fig. S3C, D) and the concomitant significant increase in PMCA (Fig. 6G, H) strongly support the hypothesis that fully differentiated effector cells acquire higher expression of PMCA to guard against apoptosis resulting from activation induced [Ca^2+^]_i_ overloads. Noteworthy is that, before cells reach full differentiation state, PMCA levels transiently decrease after short term stimulation (Fig. 6 E, F) allowing [Ca^2+^]_i_ levels required to fulfill cellular demands during initial proliferation.

CD4^+^ T cell differentiation is a complex process in which a plethora of genes are rigorously orchestrated and we hypothesized that PMCA4 is an important regulator of this process. To gain direct insight into the regulatory role of PMCA4 in CD4^+^ T cell differentiation, we monitored differentiation and compartment distribution upon activation of naïve cells while downregulating PMCA4 (siPMCA). We found that siPMCA-treated cells exhibit a skewed compartment stoichiometry following activation with more cells remaining in the naïve compartment parallel to a decreased fraction of cells able to reach the effector compartments (EM and EMRA) (Fig. 5B, C). Interestingly, although CD62L and CCR7 are often co-expressed and are commonly considered equivalent for population definition (Eichmann et al., 2020, Sallusto et al., 1999), we observed an effect of PMCA4 downregulation on the expression of CD62L (Fig. 5B, C) but not of CCR7 (Fig. S2E). This might be explained by the different kinetics of expression of both molecules were activated T cells rapidly shed CD62L of their surface and gain new set of selectins which allow them to migrate to non-lymphoid tissues (Bajnok, Ivanova et al., 2017) Moreover, a CD62L^-^CCR7^+^ population has been identified as a frequently existing memory population (Unsoeld & Pircher, 2005) and was detected in cancer studies where it has been shown to have homing and effector functions that are distinct from conventional CM and EM, though much less understood (Xie, 2013). These results implicate importance of PMCA4 in regulating early events of activation.

Our analysis of the effect of downregulation of PMCA on activation markers revealed differential effects. Transcriptional upregulation of the activation marker CD69 is triggered immediately (30-60 min) upon activation and reaches measurable levels already after 2-3h (Reddy, Eirikis et al., 2004) then declines within 6 h (Cibrian & Sanchez-Madrid, 2017). These fast kinetics could indicate that CD69 is regulated by mechanisms independent of PMCA and [Ca^2+^]_i_ and thus explain our finding that total frequency of cells expressing CD69 was not altered by siPMCA treatment (Fig. 4D, E). On the other hand, downregulation of PMCA4 resulted in significant increase of fraction of cells expressing CD154 (CD40L). CD154 belongs to the TNF gene family and its upregulation follows a biphasic pattern with the first phase being TCR dependent and taking place within the first 24 h after activation. The later phase is dependent on the cytokine composition in the cellular milieu (Lee, Haynes et al., 2002, Roy, Waldschmidt et al., 1993). Furthermore, CD154 transcription is enhanced by Ca^2+^-NFAT pathway (Daoussis, Andonopoulos et al., 2004, Wu, Chang et al., 2017) which together with an increased NFAT activity upon downregulation of PMCA4 (Boczek, Lisek et al., 2017) might explain the increase observed in CD154 expression in siPMCA-treated cells (Fig 4D, E). Together, these findings support our hypothesis that PMCA is essential for regulating differentiation processes in functional CD4^+^ T cells.

By virtue of their ability to activate, repress or modify gene expression, transcription factors regulate cell division, differentiation and function (Shi, Lee et al., 1997). Transcriptional control of PMCA isoforms has been addressed (Habib, Park et al., 2007, Ximenes, Kamagate et al., 2003) but remains to time, however, poorly understood. Here we applied *in silico* analyses (Fig. 6A) and identified YY1, a ubiquitously expressed transcription factor (Gordon, Akopyan et al., 2006), to be likely involved in modulation of PMCA4 expression. Indeed, we confirmed that YY1 binds specifically to a predicted sequence in PMCA4 promoter region (Fig. 6B). Furthermore, our results suggest a repressive effect of YY1 on PMCA expression, as indicated by the inverse correlation between expression of PMCA4 and that of YY1 over the course of differentiation (Fig. 6E-I) and more importantly, by the increased PMCA promoter activity upon downregulation of YY1 (Fig. 6J, K). There are several lines of evidence for the interaction between YY1 and c-myc (Austen, Cerni et al., 1998, Riggs, Saleque et al., 1993, Shrivastava, Yu et al., 1996). c-Myc inhibits the DNA binding capacity of YY1 (Shrivastava et al., 1996) in a direct, binding-independent (Austen et al., 1998) but c-Myc concentration dependent manner (Shrivastava et al., 1996). Moreover, Habib and co-workers have shown that in B-cells c-Myc regulates differentiation and function by mechanisms including downregulation of PMCA (Habib et al., 2007). Therefore, we hypothesize that PMCA4 is regulated by a complex of transcription factors including c-Myc and YY1 (Zeller et al., 2006, Habib et al., 2007). If and how individual members of this complex have reciprocal effects needs further investigation. By regulating PMCA4 levels and thereby shaping Ca^2+^ signals, the activities of calcineurin and NFAT, and thus T cell differentiation are modulated. With an upregulation approach, importance of PMCA to clear near plasma membrane Ca^2+^ thus allowing for NFAT activation was also demonstrated (Go, Hooper et al., 2019). Our results suggest a scenario in which another layer of transcriptional control regulates T cell differentiation and activity in which upstream to the PMCA-Calcineurn/NFAT line, expression of PMCA is regulated by YY1/c-Myc.

Together, our data suggest a model in which PMCA4 plays a pivotal role shaping Ca^2+^ signals in different activation states of CD4^+^ T cells. This role is under a biphasic transcriptional regulation by YY1. The first phase starts once CD4^+^ T cell are activated where YY1 is upregulated and represses PMCA expression so that CD4^+^ T cells can accommodate higher cytosolic [Ca^2+^]_i_ levels to support gene expression and proliferation. In the second phase, the initial repression by YY1 is alleviated and PMCA expression increases to guard against [Ca^2+^]_i_ overloads and promote survival of effector cells and formation of long lasting memory cells that are able to efficiently mediate faster immune response upon subsequent antigen encounter.

## Materials and Methods

### *T-cell isolation and* in vitro *polarization*

Blood samples were collected from healthy donors in the Institute of Clinical Hemostaseology and Transfusion Medicine, Saarland University, Homburg. Research was approved by the local ethical committee (83/15; FOR2289-TP6, Niemeyer/Alansary) and blood donors provided their written consent. Following thrombocytes apheresis, peripheral blood mononuclear cells (PBMCs) were isolated from leukocytes reduction chambers (LRS, Trima Accel^®^) according (Knorck, Marx et al., 2018, Neron, Thibault et al., 2007). Lymphocytes where enriched by overnight incubation of PBMCs to remove adherent monocyte and macrophages. On the next day CD4^+^ cells were stained and naïve (CD4^+^CD127^high^CD25^-^CD45RO^-^) or memory (CD4^+^CD127^high^CD25^-^CD45RO^+^) cells were sorted using FACSAria III (BD). Isolated CD4^+^ cells were seeded at density of 2×10^6^ cells/ml in AIMV medium containing 10% FCS and 1% Pen/Strep and cultured for at least 24 h before conducting experiments. Where indicated, naïve cells were polarized *in vitro* into Th1 or Treg according to (Kircher et al., 2018). Alternatively, where indicated, total CD4^+^ T cells were obtained from PBMC using negative isolation Kit (Miltenyi, #130-096-533) and automated cell isolation system (Mitenyi, AutoMACS).

### Transfection

For downregulation of PMCA, 2×10^5^ cells were seeded in 96-well plate in Accell siRNA Delivery Media (ADM, Horizon) supplemented with 10ng/mL of recombinant IL-2 and transfected with 1 μM accel siRNA targeting ATP2B4 (Accell Human ATP2B4 siRNA - SMARTpool, E-006118-00-0005, Horizon) or non-silencing control RNA (Accell Non-targeting Pool, D-001910-10-05, Horizon) according to the manufacturer’s instructions. Seventy-two hours later cells were harvested for mRNA or functional analysis.

### Flow cytometric analysis

For flow cytometric analysis cells were harvested, washed then stained with a viability dye (Zombie Aqua Fixable Viability Kit, Biolegend) followed by surface staining. Antibodies used for staining were supplied from Biolegend and are listed in Supplemental table 2. Flow cytometric analysis was performed with FACSVerse (BD).

### Single cell Ca^2+^ imaging

T cells were loaded in suspension with 1μM Fura 2-AM in AIMV medium at room temperature for 20-25 minutes and seeded on poly-ornithine coated glass coverslips. All experiments were at room temperature as in (Kircher et al., 2018). For stimulation of store operated Ca^2+^ influx either 1 μM Thapsigargin (Tg). Where indicated cells were treated 5 μM or 20 μM caloxin1 C2. Analyzed parameters include: maximum [Ca^2+^]_i_ (Peak), steady state [Ca^2+^]_i_ analyzed as the average [Ca^2+^]_i_ 25 s before removal of [Ca^2+^]_o_ (Plateau) and the fraction of retained [Ca^2+^]_i_ (Plateau/Peak) calculated for each cell independently. Analysis of the efflux rates were performed using Origin software. To test the influence of [Ca^2+^]_i_ on PMCA activity, cells were binned according to their [Ca^2+^]_i_ (iso-cells) and there corresponding efflux rates were compared.

### Transcription factors binding site (TFBS) analysis

A 5000 bp long sequence upstream to start codon of *ATP2B4* was used for as a potential promoter sequence for *ATP2B4*. Prediction of transcription factor potentially regulating PMCA was done using the algorithms provided by three different data bases: ChIP-Seq (Ambrosini et al., 2016), HOCOMOCO (Kulakovskiy et al., 2018) and Promo V3 (Messeguer et al., 2002). Identification of transcription factor binding site (TFBS) was done by Emsembl Trancription Factor tool (Enembl Regulatory Build (Zerbino, Wilder et al., 2015)), and making use of the matrix-based transcription factor binding site in JASPAR database (Fornes, Castro-Mondragon et al., 2020). Results obtained from the independent data bases were subject to venn analysis using the online tool Venny 2.1 (Oliveros, 2007) to detect common results, listed in supplementary table 1.

### Chromatin Immunoprecipitation (ChIP)

To test binding of transcription factors to PMCA4b promoter region, antibodies against transcription factors (YY1 or NFAT, cell signaling, 3 μg each) or isotype control antibodies (rabbit IgG or mouse IgG, respectively) were coupled to protein A/G agarose beads (Santa Cruz) in coupling buffer (0.01M sodium phosphate, 0.15M NaCl, pH 7.2) overnight at 4°C then cross-linked to the beads using 5 mM BS3 (Thermoscientific). In parallel, human CD4^+^ T cells were harvested, washed then cross-linked with PBS then containing 5 mM DSP (Thermoscientific) and 3% PFA for 20 min at 4°C. Excess crosslinker and PFA were quenched by 125 mM Glycine in PBS. Cell pellets were lysed in 20mM Tris, 100mM KCl, 10% glycerine (v/v), 0.5% n-Octyl-β-D-glucopyranoside (Merck Millipore). Finally lysates were subjected to chromatin shearing with Qsonica Sonicator Q700 (Thermoscientific) in total processing time of 4 min, 20 s laps of 40% amplitude and 10 s pauses. Clear lysates were diluted to 1 ml with binding buffer (20mM Tris, 100mM KCl, 10% glycerine (v/v)). An aliquot of 50 μl was kept as an input control. Lysates were incubated with preclearing beads (isotype IgG bound beads) for 3 h at 4°C, unbound lysate was allowed to bind to control IgG (a fresh aliquot of isotype IgG bound beads) for 3 h at 4°C and finally unbound lysate was bound to chromatin IP beads (transcription factor bound beads) for 3 h at 4°C. Beads collected at different steps were washed with standard low salt, high salt, LiCl buffer and finally with TE according to Thermoscientific protocol. Finally, DNA was eluted with 1% SDS in 100 mM NaHCO3, purified using DNA purification kit (Qiagen) and used as template to test for binding of the PMCA4b promoter region using Q5 hot start polymerase (NEB biolabs) and the primers: forward 5’-CACCACCGTGCCAGCTAA-3’, reverse 3’-GAAGGGTGCTAGTTGGACA-5’. Amplicons were visualized by DNA agarose and identity was confirmed by sanger sequencing.

### Dual luciferase reporter assay

To estimate alterations in PMCA4 promoter activity, the region (Gene ID NG_029589.1, bp 663-1052) identified by ChIP assay to be potentially bound by YY1 was amplified with similar primers to as above mentioned but adding SacI and NheI recognition sequences. The corresponding cDNA was subcloned into pGL3 basic (Promega, no. E1751) upstream to sequence encoding redfirefly luciferase. Three days before conducting the assay, HEK293 cells were transfected with siRNA targeting YY1 (siYY1) (Accel Human YY1 siRNA SMART pool, no. E-011796-00-0010, Horizon) or with non-silencing control RNA (ns). Forty-eight hours later, cells were transfected with the created plasmid (PMCA Promoter, PP) or pGL3 basic (ϕ) together with plasmid encoding green renilla luciferase under thymidine kinase promoter control (pTK-Green Renilla Luciferase, ThermoFisher Scientific, no. 16154) in a DNA ratio of 2:1. Finally, 24h later, cells were stimulated with 5ng/ml phorbol 12-myristate 13-acetate (PMA) and 500 ng/ml ionomycin for 6h then lysed in passive lysis buffer commercially provided in the dual luciferase reporter assay kit (Promega, no. E1910). Assay was conducted according to manufacturer’s instructions and luminescence measurements were done using CLARIOstar (BMG, LABTEC) at gain of 3600. For analysis, red firefly luciferase luminescence was normalized to the green renilla luciferase and data presented as average ± SEM of the relative change to the control condition of each independent experiment.

### RNA isolation, cDNA synthesis, and quantitative real-time PCR (qRT-PCR)

The indicated cell types were harvested and stored at −80°C until RNA was isolated using RNeasy kit (Qiagen) following manufacturer’s instructions. SuperScriptTMII Reverse Transcriptase (Life technologies) was used to generate complementary DNA (cDNA) and subsequent qRT-PCR was conducted using QuantiTect SYBR Green Kit (Qiagen) and a CFX96 Real-Time System (Biorad). For quantification, threshold cycle (Cq) values of a gene of interest was normalized to that of either RNA polymerase or to TATA box binding protein (TBP) using the ΔCq method. For comparison of mRNA levels between two or more populations, fold change was calculated with 2^-ΔΔCq^ method.

### Westerm Blot

For protein expression analysis cells were harvested, washed with PBS then lysed in buffer containing 20mM Tris, 100mM KCl, 10% glycerine (v/v), 0.5% n-Dodecyl beta Maltoside (DDM), pH 7.4. Lysates were stored at −80° until analyzed. Standard SDS-PAGE was performed followed by electrotransfer to nitrocellulose membranes. Immunoblots were probed with anti-Orai1 (Sigma, catalogue number O8264), anti-STIM1 (Proteintech, # 11565-1-A), anti-STIM2 (Sigma, # S8572), anti-GAPDH (Cell Signalin, clone 14C10) anti-ß-actin (Abcam, clone AC-15) or anti-PMCA4 (Thermoscientific, clone JA9) at a 1:1000 dilution. For protein detection an enhanced chemiluminescence detection reagent was used (Clarity Western ECL Substrate, Biorad). Densitometric quantification of detected protein bands was done with Quantity one software (Biorad).

### Statistical analysis

For analysis of Ca^2+^ imaging data, 100-250 cells were measured per experiment, 1-2 independent experiments were done per donor from up to 14 donors. Otherwise, data is presented as mean ± S.E.M. Data was tested for normal distribution. When comparing two groups statistical significance was tested by performing unpaired, two-tailed Student Student t-test for normally distributed data sets and Mann-Whitney test when samples are not normally distributed or when the sample size was not sufficient to test for normality. Comparing more than two groups was done using Kruskall-Wallis one-way analysis of variance (ANOVA) or conventional ANOVA when data showed normal distribution and equal variance. Asterisks indicate significant differences for different p values as follows: * p<0.05, ** p<0.01, *** p< 0.001. Statistical analysis was performed using Graphpad Prism

## Acknowledgement

We kindly thank Dr. Eichler and the Institute for Clinical Hemostaseology and Transfusion Medicine for providing donor blood; Dr. Schwarz for supervising primary cell isolation and critical reading of MS; Dr. Hoth for technical and equipment support and critical reading of MS; Anja Bergsträßer and Dr. Krause from the FACS Facility of the Institute of Physiology (DFG grant 207087572) for help with cell sorting; Carmen Hässig, and Kathrin Förderer for technical assistance and Dr. Martin Hart and Tim Kehl for help with the TFBS analysis, Dr. Prates-Roma and Dr. Frisch for help with promoter assays. Our work is supported by the Deutsche Forschungsgemeinschaft DFG (FOR 2289-P6) to B.A.N and D.A. and (TRR219-C09) to B.A.N. The FACSverse was funded by DFG (GZ: INST 256/423-1 FUGG).

## Author Contributions

All authors conceptualized the study and edited the manuscript, M.M.W, and D.A. performed experiments and data analysis, B.A.N. and D.A. secured funding and resources, D.A. wrote the manuscript.

## Declaration of Interests

The authors declare no competing interests.

## Supplementary Figure Legends

**Supplementary Figure 1:**
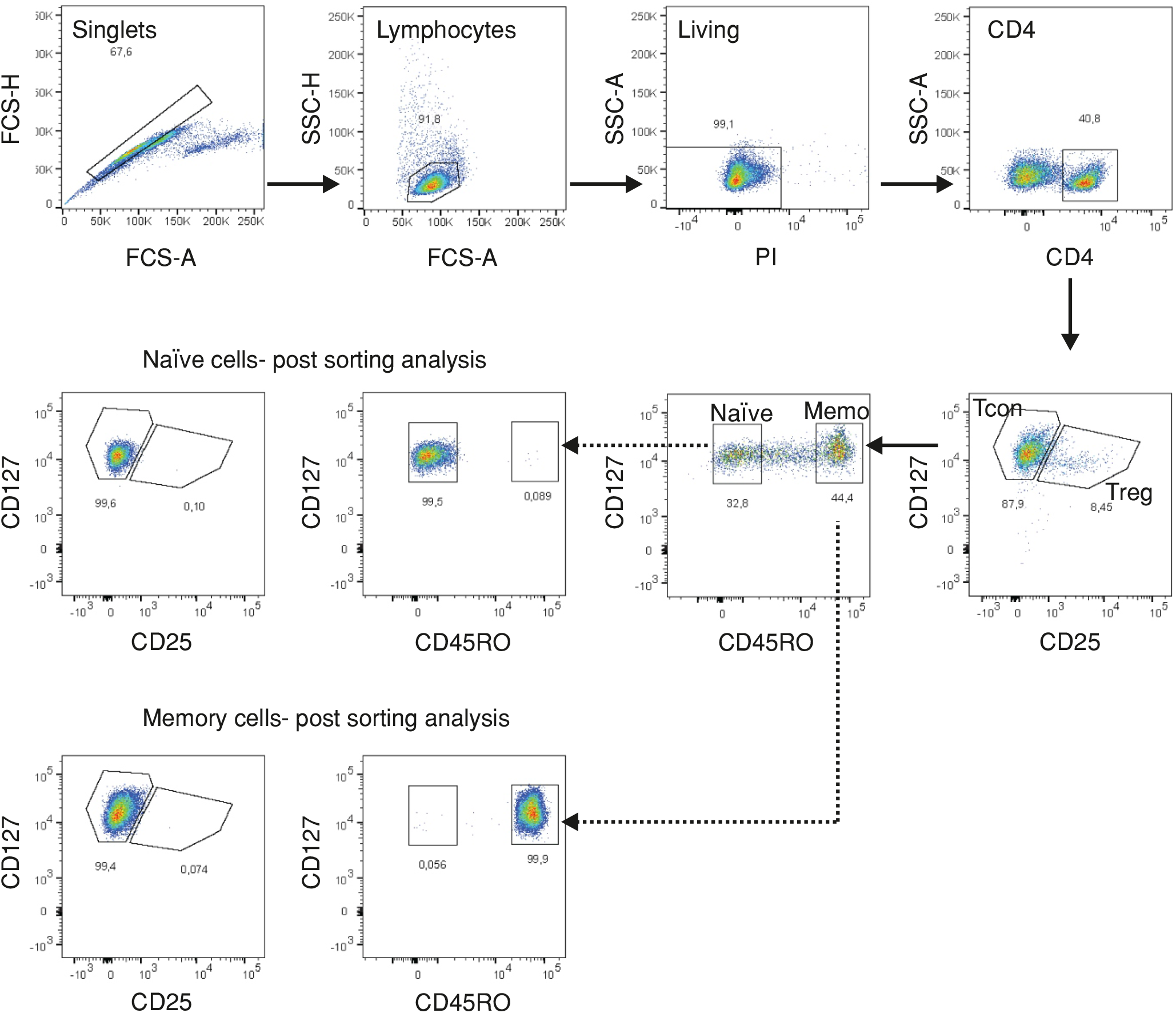
Sorting strategy of naïve and memory cells. Representative flow cytometry images showing the isolation FACSorting strategy of human naïve (CD4^+^CD127^high^CD25^-^CD45RO^-^) or memory (CD4^+^CD127^high^CD25^-^CD45RO^+^) T cells using FACSAria III (BD).

**Supplementary Figure 2:**
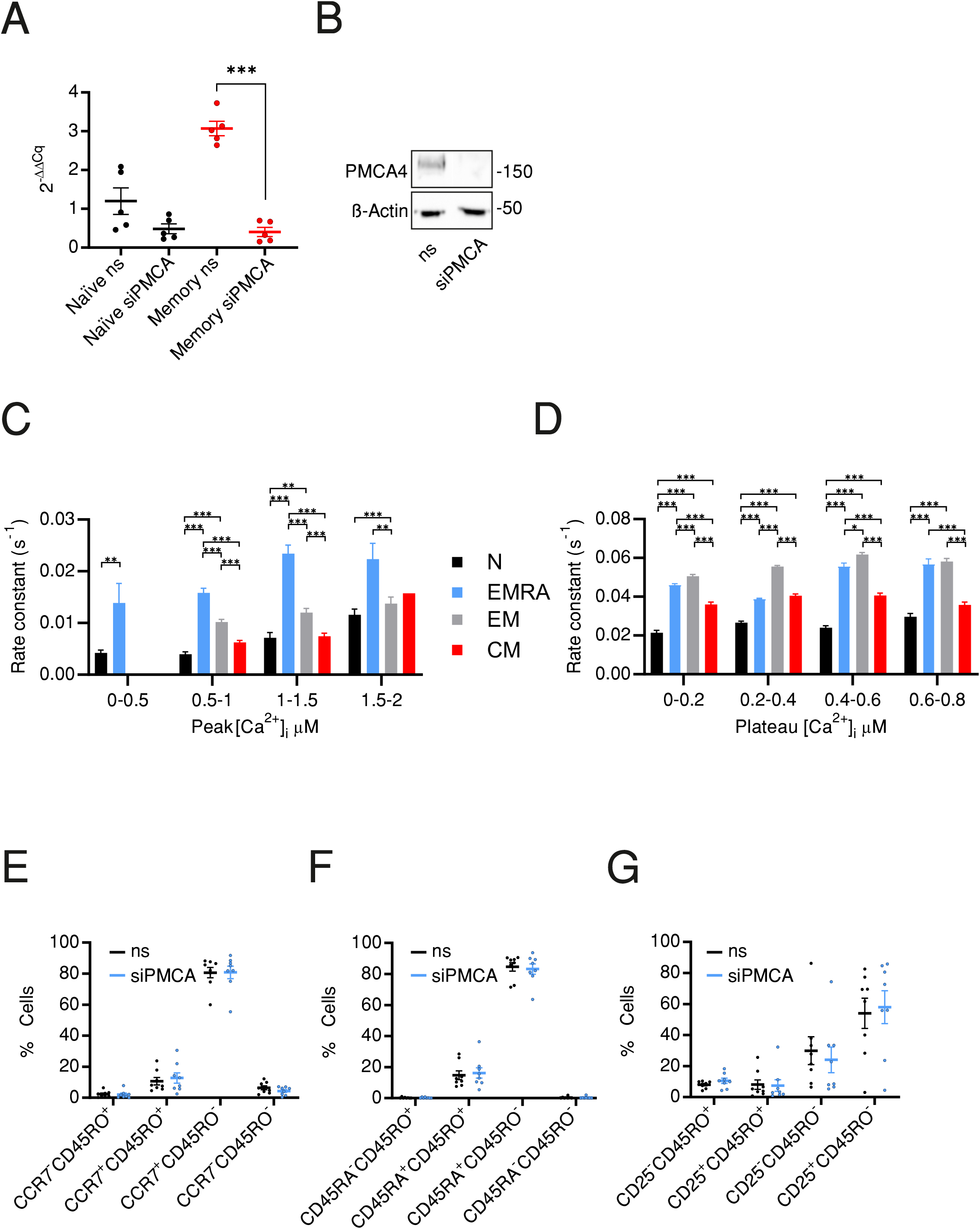
siRNA-mediated downregulation of PMCA and Ca^2+^ efflux rates of cellular compartments. (A) Quantitative RT-PCR analysis showing relative expression level of PMCA4 in naïve (black) and memory (red) cells transfected with non-silencing RNA (ns) or with siRNA targeting PMCA4 (siPMCA). (B) Representative WB showing PMCA4 expression in total CD4 T cells transfected with non-silencing RNA (ns) or with siRNA targeting PMCA4 (siPMCA). (C) Analysis of Ca^2+^ efflux rates in presence of [Ca^2+^]_o_ in cells measured in Fig. 4E as function of peak [Ca^2+^]_i_ (D) Analysis of Ca^2+^ efflux rates after removal of [Ca^2+^]_o_ in cells measured in in Fig. 4F as function of plateau [Ca^2+^]_i_. (E-G) Average fractions of cells expressing the indicated surface markers as obtained by flow cytometry analysis of naïve cells transfected with non-silencing RNA (ns, black) or with siRNA targeting PMCA4 (siPMCA, blue) for 72 h then stimulated with antiCD3/CD28 coated beads for further 24 h as in Fig.5.

**Supplementary Figure 3:**
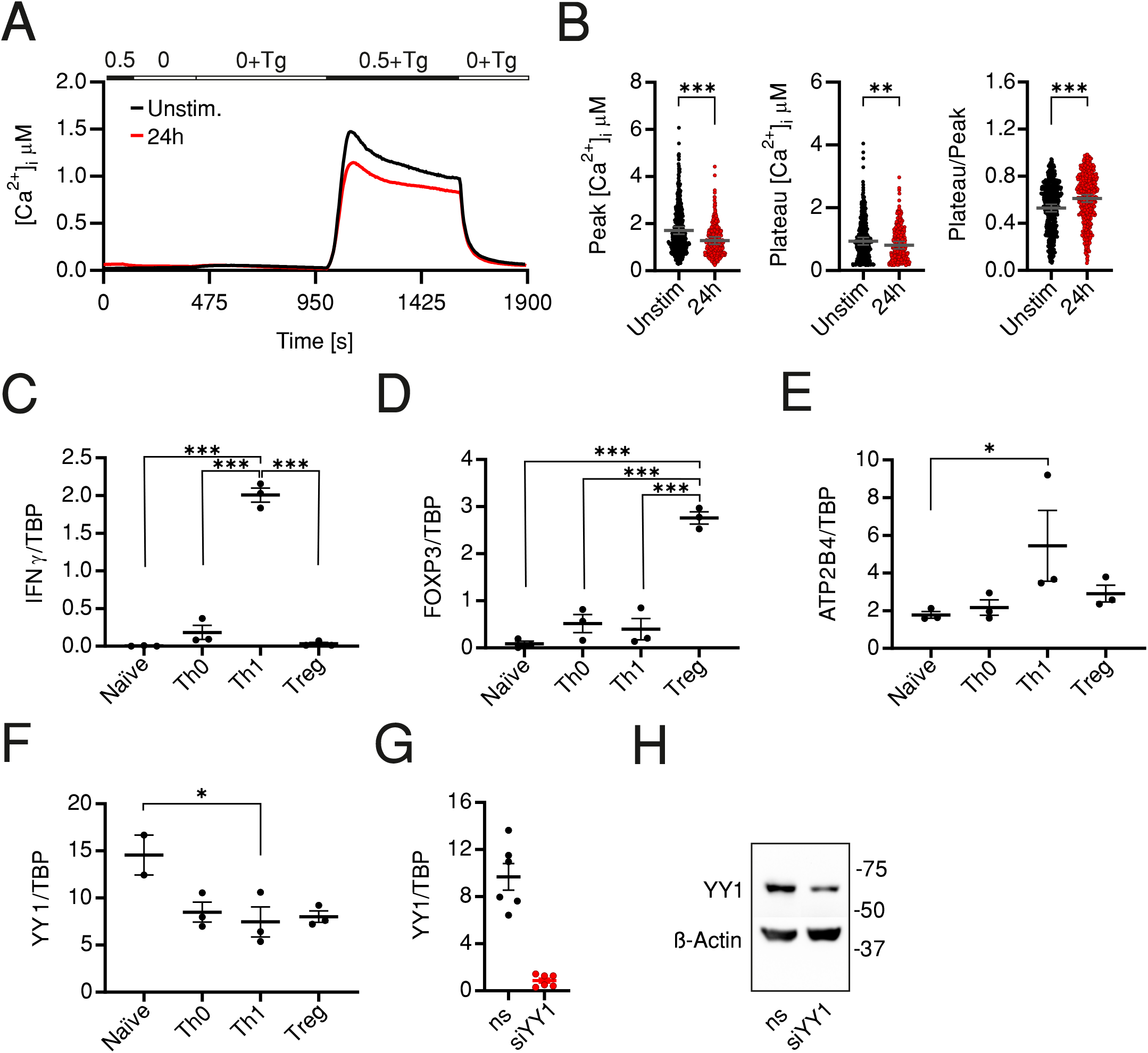
SOCE profile and expression analysis of genes of interest in activated cells. (A) Average traces showing changes of [Ca^2+^]_i_ over time in response to changes of extracellular solutions containing the shown [Ca^2+^]_o_ indicated in the bar above the traces. SOCE was measured in resting total human CD4^+^ cells (Unstim., black) or following a 24h antiCD3/CD28 coated bead stimulation (24h, red). (B) Bar graphs showing analyzed parameters of SOCE measured in (A). (C-E) Quantitative RT-PCR analysis of IFNG (C), FOXP3 (D), ATP2B4 (E) or YY1 (F) measured in resting naive CD4^+^ T cells (naïve) or cells cultured for 7 d under control conditions (Th0) or conditions to polarize cells into Th1 or Treg. (G) Quantitative RT-PCR analysis showing relative expression level of YY1 in total CD4^+^ T cells transfected with non-silencing RNA (ns) or with siRNA targeting YY1 (siYY1). (B) Representative WB showing YY1 expression in total CD4^+^ T cells transfected with non-silencing RNA (ns) or with siRNA targeting YY1 (siYY1).

**Supplemental table 1:**
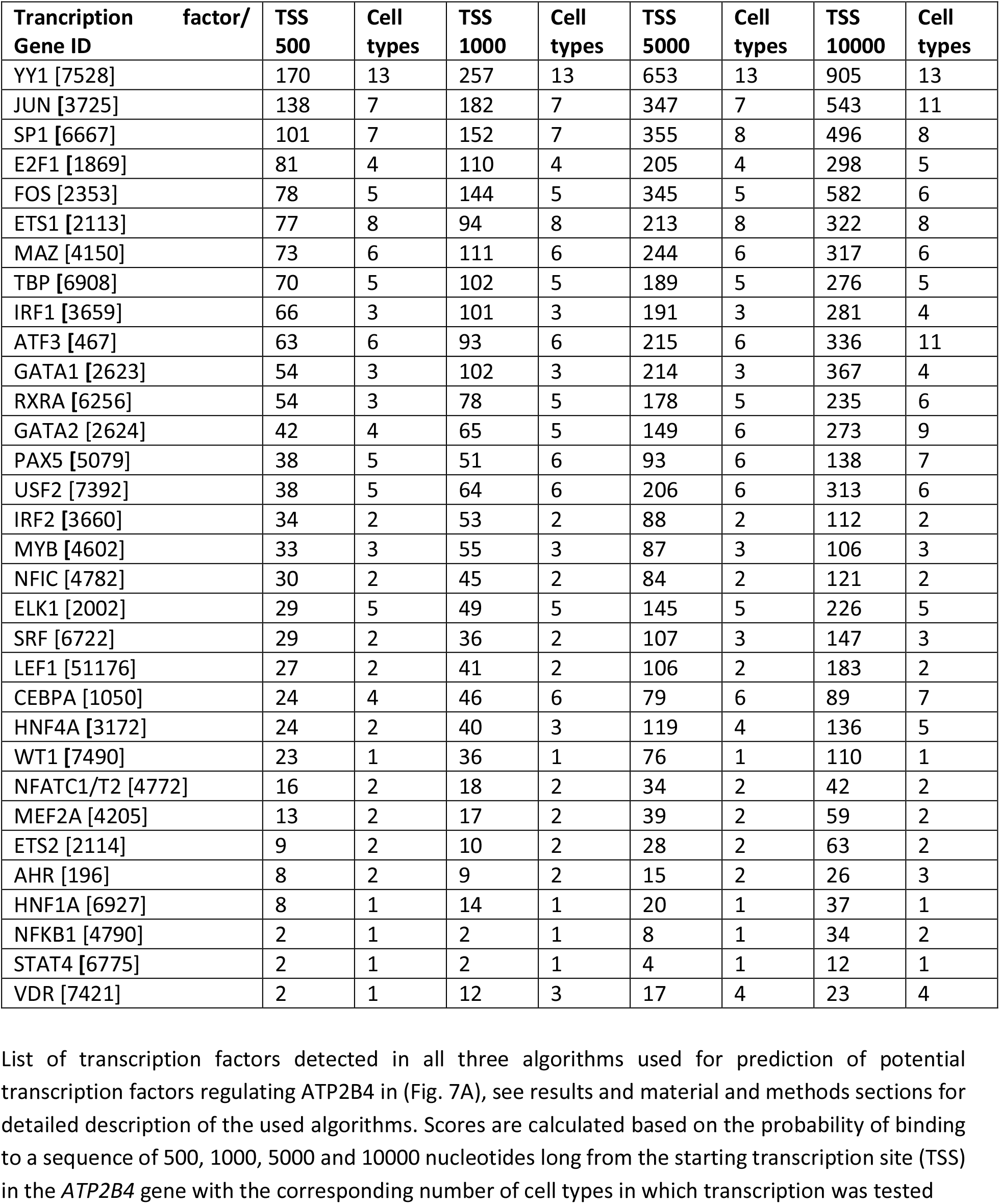
List of transcription factors identified by in silico analysis

| Transcription factor/<br>Gene ID | TSS<br>500 | Cell<br>types | TSS<br>1000 | Cell<br>types | TSS<br>5000 | Cell<br>types | TSS<br>10000 | Cell<br>types |
| --- | --- | --- | --- | --- | --- | --- | --- | --- |
| YY1 [7528] | 170 | 13 | 257 | 13 | 653 | 13 | 905 | 13 |
| JUN [3725] | 138 | 7 | 182 | 7 | 347 | 7 | 543 | 11 |
| SP1 [6667] | 101 | 7 | 152 | 7 | 355 | 8 | 496 | 8 |
| E2F1 [1869] | 81 | 4 | 110 | 4 | 205 | 4 | 298 | 5 |
| FOS [2353] | 78 | 5 | 144 | 5 | 345 | 5 | 582 | 6 |
| ETS1 [2113] | 77 | 8 | 94 | 8 | 213 | 8 | 322 | 8 |
| MAZ [4150] | 73 | 6 | 111 | 6 | 244 | 6 | 317 | 6 |
| TBP [6908] | 70 | 5 | 102 | 5 | 189 | 5 | 276 | 5 |
| IRF1 [3659] | 66 | 3 | 101 | 3 | 191 | 3 | 281 | 4 |
| ATF3 [467] | 63 | 6 | 93 | 6 | 215 | 6 | 336 | 11 |
| GATA1 [2623] | 54 | 3 | 102 | 3 | 214 | 3 | 367 | 4 |
| RXRA [6256] | 54 | 3 | 78 | 5 | 178 | 5 | 235 | 6 |
| GATA2 [2624] | 42 | 4 | 65 | 5 | 149 | 6 | 273 | 9 |
| PAX5 [5079] | 38 | 5 | 51 | 6 | 93 | 6 | 138 | 7 |
| USF2 [7392] | 38 | 5 | 64 | 6 | 206 | 6 | 313 | 6 |
| IRF2 [3660] | 34 | 2 | 53 | 2 | 88 | 2 | 112 | 2 |
| MYB [4602] | 33 | 3 | 55 | 3 | 87 | 3 | 106 | 3 |
| NFIC [4782] | 30 | 2 | 45 | 2 | 84 | 2 | 121 | 2 |
| ELK1 [2002] | 29 | 5 | 49 | 5 | 145 | 5 | 226 | 5 |
| SRF [6722] | 29 | 2 | 36 | 2 | 107 | 3 | 147 | 3 |
| LEF1 [51176] | 27 | 2 | 41 | 2 | 106 | 2 | 183 | 2 |
| CEBPA [1050] | 24 | 4 | 46 | 6 | 79 | 6 | 89 | 7 |
| HNF4A [3172] | 24 | 2 | 40 | 3 | 119 | 4 | 136 | 5 |
| WT1 [7490] | 23 | 1 | 36 | 1 | 76 | 1 | 110 | 1 |
| NFATC1/T2 [4772] | 16 | 2 | 18 | 2 | 34 | 2 | 42 | 2 |
| MEF2A [4205] | 13 | 2 | 17 | 2 | 39 | 2 | 59 | 2 |
| ETS2 [2114] | 9 | 2 | 10 | 2 | 28 | 2 | 63 | 2 |
| AHR [196] | 8 | 2 | 9 | 2 | 15 | 2 | 26 | 3 |
| HNF1A [6927] | 8 | 1 | 14 | 1 | 20 | 1 | 37 | 1 |
| NFKB1 [4790] | 2 | 1 | 2 | 1 | 8 | 1 | 34 | 2 |
| STAT4 [6775] | 2 | 1 | 2 | 1 | 4 | 1 | 12 | 1 |
| VDR [7421] | 2 | 1 | 12 | 3 | 17 | 4 | 23 | 4 |

**Supplemental Table 2:**
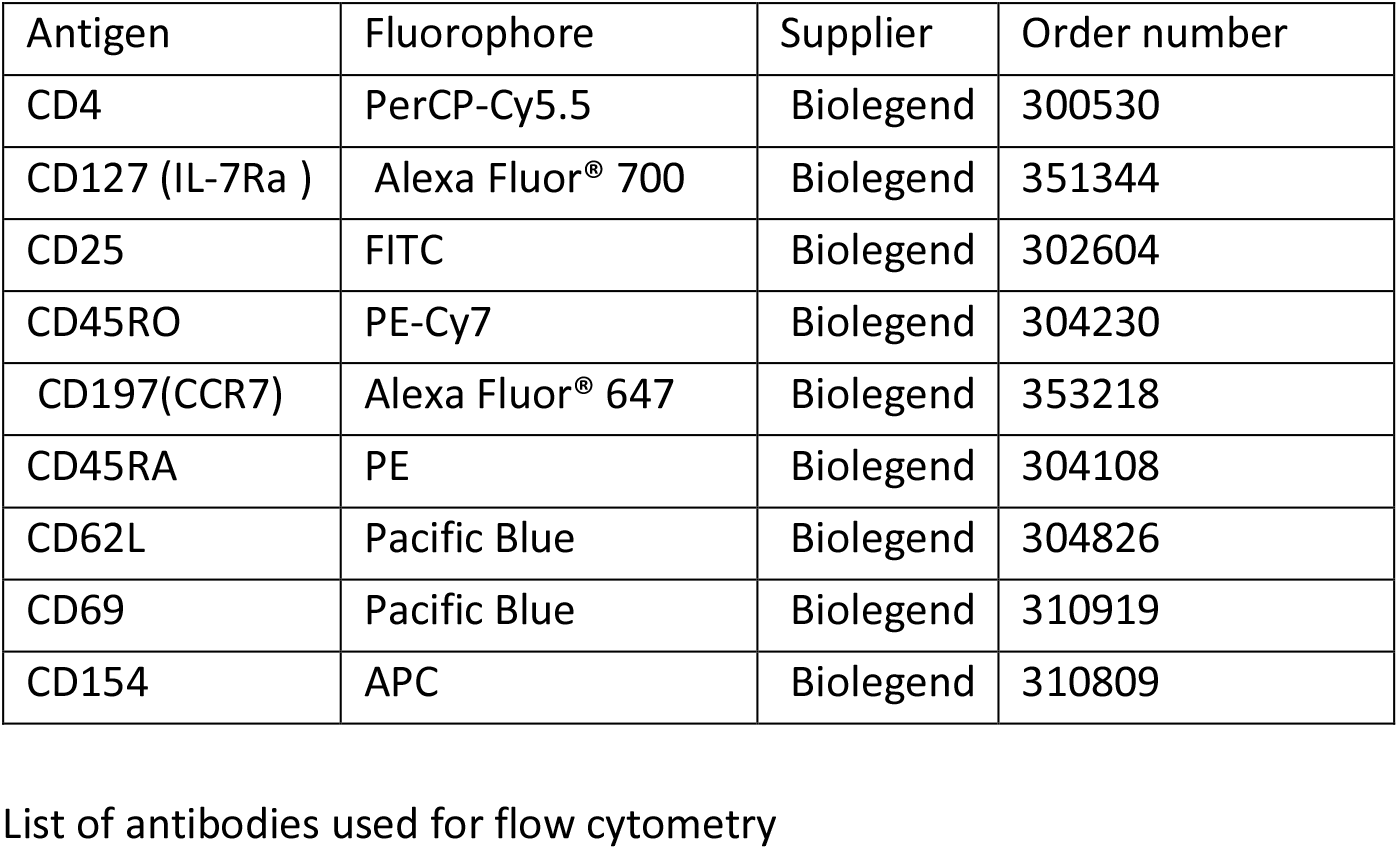
List of antibodies used for flow cytometry

